# How does literacy affect speech processing? Not by enhancing cortical responses to speech, but by promoting connectivity of acoustic-phonetic and graphomotor cortices

**DOI:** 10.1101/2021.06.04.446930

**Authors:** Alexis Hervais-Adelman, Uttam Kumar, Ramesh K. Mishra, Viveka N. Tripathi, Anupam Guleria, Jay P. Singh, Falk Huettig

**Affiliations:** Neurolinguistics, University of Zurich, Department of Psychology, Binzmühlerstrasse 14, 8050, Zurich, Switzerland; Neuroscience Center Zurich, University of Zurich and Eidgenössische Technische Hochschule Zurich, 8057 Zurich, Switzerland; Centre of Biomedical Research, Raibareli Road, Lucknow-226014, Uttar Pradesh, India; University of Hyderabad, Prof. C.R. Rao Road, Gachibowli, Hyderabad-500046, Telangana, India; Centre for Behavioural and Cognitive Sciences, University of Allahabad, University Road, Old Katra, Allahabad-211002, Uttar Pradesh, India; Max Planck Institute for Psycholinguistics, Wundtlaan 1, 6525 XD Nijmegen, The Netherlands

## Abstract

Previous research suggest that literacy, specifically learning alphabetic letter-to-phoneme mappings, modifies online speech processing, and enhances brain responses to speech in auditory areas associated with phonological processing (Dehaene et al., 2010). However, alphabets are not the only orthographic systems in use in the world, and hundreds of millions of individuals speak languages that are not written using alphabets. In order to make claims that literacy *per se* has broad and general consequences for brain responses to speech, one must seek confirmatory evidence from non-alphabetic literacy. To this end, we conducted a longitudinal fMRI study in India probing the effect of literacy in Devanagari, an abubgida, on functional connectivity and cerebral responses to speech in 91 variously literate Hindi-speaking individuals. Twenty-two completely illiterate participants underwent six months of reading and writing training. Devanagari literacy increases functional connectivity between acoustic-phonetic and graphomotor brain areas, but we find no evidence that literacy changes the way speech is processed, either in cross-sectional or longitudinal analyses. These findings shows that a radical reconfiguration of the neurofunctional substrates of online speech processing is not a universal result of learning to read, and raise the possibility that writing, not only reading, may be instrumental in moulding literate speech perception.

**Significance Statement:** It has come to be accepted that a consequence of being able to read is enhanced auditory processing of speech, reflected by increased cortical responses in areas associated with phonological processing. Here we find no relationship between literacy and the magnitude of brain response to speech stimuli in individuals who speak Hindi, which is written using a non-alphabetic script, Devanagari - an abugida. We propose that the exact nature of the script under examination must be considered before making sweeping claims about the consequences of literacy for the brain. Further, we find evidence that literacy enhances functional connectivity between auditory processing areas and graphomotor areas, suggesting a mechanism whereby learning to write, not only to read, might influence speech perception.

## Introduction

Learning to read and write involves acquiring a mapping of spoken language to orthographic symbols, which have both a visual (recognition) and a motor (production) component. These mapping processes have been suggested to have functional consequences for brain areas associated with acoustic-phonetic processing that are manifest during the processing of speech. It has been argued that literacy modifies the underlying phonological code for speech reception, as evidenced in increased brain responses to speech in auditory areas in literate compared to illiterate individuals (Dehaene et al., 2010). But how universal are such effects? While literacy in *alphabetic* script may induce such changes, alphabetic scripts encode subsyllabic segments, many of which cannot be produced in isolation (i.e. most consonants) and may therefore incur different phonological restructuring demands to other types of scripts that encode producible speech units (e.g. syllabaries, logosyllabaries and abugidas, Daniels, 2020). The worldwide diversity of orthographic systems invites closer examination of these findings to verify whether they hold for non-alphabetic writing systems. In psychology the pitfalls of drawing conclusions for the whole of humanity based only on the so-called “WEIRD” (Western, educated, industrialised, rich, democratic) populations (Henrich, Heine, & Norenzayan, 2010) are increasingly understood, and the study of literacy is no exception. In this study, we aimed to extend the generalizability of conclusions regarding the impact of literacy on brain responses to speech.

Notwithstanding the particularities of the orthographic system at hand, literacy is almost never acquired as a purely receptive skill, but typically involves a significant motor component when learning to write. Nevertheless, the role of learning to write in developing acoustic-phonetic representations at the level encoded by the script is barely discussed, even though there would be every reason to posit that creating motor-auditory mappings may lead to adaptations analogous to those of learning the visual-auditory mappings of script.

In this study, we used fMRI to record cerebral responses to auditory sentences in a group of 91 Hindi-speaking individuals of varying levels of literacy (ranging from illiterate to fluent readers) from rural communities in Northern India, first in cross-sectional investigation, and in a follow-up investigation after 22 illiterate participants had undergone a six-month long literacy training intervention. We deliberately used a listening task with no meta-linguistic component to ensure that any observed effects were not the result of deliberate, task-driven, phonological processing components. We also examined functional connectivity to phonological processing regions during speech processing in order to evaluate the potential visuo-auditory and graphomotor-auditory links that may be modified in literate individuals. Analyses were carried out to seek a relationship between literacy and brain response, at the whole-brain level and in three regions of interest: the left Planum Temporale (PT) and left Posterior Superior Temporal Gyrus (pSTG) both associated with acoustic-phonetic processing of speech and the Visual Word-Form Area (VWFA, an area of left posterior fusiform gyrus with a particular sensitivity to orthographic stimuli).

Contrary to the assumption that literacy necessarily induces changes to online speech processing, we find no evidence for changes to online speech processing, either cross-sectionally or longitudinally, with Bayesian analyses suggesting substantial evidence in favour of there being no effect. However, we find significant evidence of literacy-enhanced functional connectivity between acoustic-phonetic processing areas of posterior superior temporal cortex with graphomotor areas, which is suggestive of the development of a functional link between a graphemic, as well as visual, code and speech processing. The findings call into question the widely-accepted impact of literacy on speech processing and also invite closer examination of the role of writing, not just reading, in shaping acoustic-phonetic processing of speech.

## Materials and Methods

The data presented in this paper includes a novel reanalysis of a subset of previously published data (Hervais-Adelman et al., 2019).

### Participants

Participants were recruited from two villages near the city of Lucknow in the Northern Indian state of Uttar Pradesh as part of a study that was approved by the ethics committee of the Center of Biomedical Research, Lucknow. After giving informed consent, 91 healthy right-handed human volunteers without a known history of psychiatric disease or neurological condition took part in the study (see Demographic and Behavioural Data for more details). All of the participants were right-handed and examined by a medical doctor. None of the participants had any known neurological impairments. Participants were interviewed about their educational background. A word-reading test and a letter identification task were administered.

### Participant Characteristics

Information on age, income and number of literate family members was obtained in an interview. Right-handedness was verified in interview by asking participants which hand they used for common activities (e.g. drawing). Raven’s Progressive Matrices were administered to test non-verbal cognitive abilities. Two measures of literacy were recorded, namely letter identification (ability to name the 46 primary Devanagari characters, called “Akshara”) and word reading ability (ability to read out-loud 86 words of varying syllabic complexity). In the akshara-identification task participants received a recorded spoken instruction. In the spoken instruction, they were told that they would be presented with the letters of the Hindi alphabet one by one. They were told that each letter would be shown for five seconds followed by a question mark. The instruction specified that when the question mark was on the screen they should name the letter out loud. The response was recorded. The recording terminated automatically after ten seconds. The 46 akshara of Devanagari were presented in font Mangal (size 96 point). The word reading test consisted of 86 words, presented one-at-a-time in font Mangal (size 96 point). The words were of differing syllabic complexity (26 monosyllabic, 30 disyllabic, and 30 trisyllabic) and presented in a pseudorandomized order. Each word was displayed on the screen for ten seconds followed by a question mark. A spoken recorded instruction specified that when the question mark was on the screen participants should say aloud the word they had seen. Participants’ responses were recorded. The recording terminated automatically after thirty seconds. Participant characteristics and correlations between reading scores and other factors are shown in Table 1, pairwise relationships between participant characteristics are shown in Table 2.

**Table 1:** Participant Characteristics, Time 1. The mean values for the N=91 participants included at Time 1. Relationships between each variable and Word Reading was calculated using non-parametric correlation (Kendall’s ⊤ -b), Significance flags: p-values: **<.01, ***<.001

| | Mean | Std. Deviation | Minimum | Maximum | Relationship with word reading scores<br>Kendall's $\tau$ |
| --- | --- | --- | --- | --- | --- |
| **Age (years)** | 28.165 | 5.57 | 18 | 40 | -0.214 |
| **Monthly income (IRN)** | 2109.89 | 1277.589 | 0 | 9000 | 0.119 |
| **N Family members** | 4.506 | 1.525 | 2 | 8 | -0.02 |
| **N Literate Members** | 2.275 | 1.434 | 0 | 6 | -0.086 |
| **Years of Schooling** | 2.22 | 3.47 | 0 | 12 | 0.605*** |
| **Word reading (max. 86)** | 25.538 | 33.301 | 0 | 86 | n/a |
| **Akshara Identification (max. 46)** | 21.132 | 17.458 | 0 | 46 | 0.746*** |
| **Raven's Matrices** | 15.385 | 4.997 | 8 | 35 | 0.242** |

**Table 2:** Pairwise Correlations Between Biographical Factors.

| Variable | Word reading | Akshara Identification | Years of Schooling | Raven's Matrices | Age | Sex (dummy coded: F=0, M=1) | Monthly Income | N. Family Members |
| --- | --- | --- | --- | --- | --- | --- | --- | --- |
| Word reading | — |  |  |  |  |  |  |  |
| Akshara Identification | 0.746, p<.001 | — |  |  |  |  |  |  |
| Years of Schooling | 0.605, p<.001 | 0.58, p<.001 | — |  |  |  |  |  |
| Raven's Matrices | 0.242, p=.002 | 0.278, p<.001 | 0.239, p=.005 | — |  |  |  |  |
| Age | -0.214, p=.006 | -0.228, p=.003 | -0.383, p<.001 | -0.088, p=.25 | — |  |  |  |
| Sex (dummy coded: F=0, M=1) | 0.42, p<.001 | 0.411, p<.001 | 0.491, p<.001 | 0.207, p=.021 | -0.209, p=.02 | — |  |  |
| Monthly Income | 0.119, p=.147 | 0.108, p=.171 | 0.043, p=.626 | -0.064, p=.423 | 0.061, p=.446 | 0.039, p=.677 | — |  |
| N. Family Members | -0.02,<br>p=.817 | -0.012,<br>p=.888 | -0.063,<br>p=.495 | -0.149,<br>p=.072 | 0.287,<br>p<.001 | -0.037,<br>p=.701 | -0.006,<br>p=.943 | — |
| N. Literate Family Members | -0.086,<br>p=.367 | -0.058,<br>p=.527 | -0.101,<br>p=.327 | -0.229,<br>p=.014 | 0.151,<br>p.104 | -0.036,<br>p=.741 | -0.145,<br>p=.139 | 0.558,<br>p<.001 |

There was an imbalance in the sex of the participants as a function of literacy status, with illiterate participants being more likely to be female. This is driven by cultural factors at the study site - literate participants acquired literacy during formal education, and there has historically been a bias towards male children receiving formal schooling. The potential consequences of this are not readily predictable, though, as discussed in a previous report on these data (Hervais-Adelman et al., 2019) existing findings do not suggest that the reading network is systematically differently-localized in female compared to male samples. We therefore do not believe that this limitation would account for effects that we observe.

### MRI Procedure

Stimuli were presented blocked by condition. There were ten blocks for each task, which were arranged in a different pseudo-random order for each participant. Stimulus presentation was controlled using E-Prime (Psychology Software Tools).

#### Stimuli

Six stimulus categories were presented in the localiser block: Visual Sentences, Auditory Sentences, Horizontal Checkerboards and Vertical Checkerboards. Two further conditions were included that invited participant responses – visual (written) and spoken commands. These commands asked the participant to respond by pressing either the left or right response button of an MR-Compatible button box. Behavioural responses to these trials are not analysed and the conditions are not described further. Ten blocks of each stimulus were presented in a randomized order (randomized per participant). In each visual sentence block each trial consisted of a simple sentence, which was shown on four successive screens with 1-3 words on each screen. Each screen was shown for 400 ms with an interval of 100 ms between each screen. Between each sentence there was a 500 ms pause. All words were displayed in font ‘Mangal’ size 86. The words were shown at the centre of the screen. Participants received a recorded auditory instruction to read the sentences. Blocks lasted 33 seconds.

For auditory sentences, each block consisted of ten sentences. Sentences were presented auditorily with four audio sequences comprising 1-3 words for each sentence. Participants were instructed to listen to the sentences carefully. Blocks lasted approximately 60s (mean: 59.55, range: 57.75 – 66.19s).

Vertical and horizontal checkerboard blocks each consisted of 30 flashing vertical checkerboards. The checkerboards changed their contrast after 400 ms. Each block lasted 12 seconds (30×400 ms). Analyses of these stimuli are not presented here, they have previously been discussed (Hervais-Adelman et al., 2019).

### Regions of Interest

#### ROI Definitions

A goal of the investigation was to probe the claims of Dehaene and colleagues (Dehaene et al., 2010) regarding the change in responsiveness to speech in PT as a function of literacy. We therefore defined a target ROI based on the coordinates of the peak effect of literacy on response to spoken language processing in their investigation. A sphere of radius 8mm centred at MNI (x,y,z mm): −38, −28, 18.

However, since Dehaene et al.’s interpretation of their finding is at the level of phonological processing of speech, it seems important to test the possibility that other phonetic processing regions may be affected by literacy. Given recent functional imaging and intracranial reports of the role of posterior superior temporal gyrus in processing of speech, particularly at the phonological level, a second ROI was created in the left posterior superior temporal gyrus (pSTG). This ROI was based on the coordinates published by Chevillet and colleagues (Chevillet, Jiang, Rauschecker, & Riesenhuber, 2013) as a focus of acoustic-phonetic processing based on an fMRI investigation. A sphere of radius 8mm centred at MNI (x,y,z mm): −56, −50, 8 was used. This ROI was investigated in order to probe whether sites other than PT relevant to phoneme processing display any modulation of activation during as a function of literacy.

A VWFA ROI was created as a sphere of 8mm radius centred upon the coordinates reported in Hervais-Adelman et al. (2019), MNI (x,y,z mm): −45, −55, −10. This ROI constitutes a control region, where the effect of literacy on response to orthographic stimulation is already known, and serves to validate the analyses executed on the two other ROIs. Loci of the ROIs are shown in Figure 4.

#### ROI data extraction

Individual participant’s T statistic for the contrast against baseline was extracted for all voxels in the ROI (excluding any missing values, i.e. voxels not containing brain tissue), and the mean was tested for a relationship with Literacy using Kendall’s T. Analyses were conducted in MNI space, using normalised single-subject images.

### MRI data acquisition and pre-processing

Anatomical and functional data were collected before and after the literacy program using a 3.0 Tesla Siemens MAGNETOM Skyra (Siemens AG, Germany) whole body magnetic resonance scanner using a 64-channel radiofrequency head coil. T1-weighted three-dimensional magnetization-prepared rapid-acquisition gradient echo (MPRAGE) images were obtained using a pulse sequence with TR=1.690ms, TE=2.60ms, TI=1.100ms, FOV=256×256, matrix size=256×256×192 and voxel size= 1.0×1.0×1.0mm^3^. Functional images for the visual and localizer runs were acquired as continuous EPI (TR = 2400ms, TE=30ms, 38 slices, voxel size: 3.5 * 3.5 * 3mm, no interslice gap, interleaved slice order). Pre-processing was carried out using the default pipeline implemented in the Conn toolbox (Whitfield-Gabrieli & Nieto-Castanon, 2012), version CONN20.b, SPM12 build 7219. This involves functional realignment and unwarping, slice-timing correction (for Siemens interleaved acquisitions), both using SPM12 default settings, followed by outlier identification based on the observed BOLD signal and subject motion parameters. Acquisitions with frame-by-frame displacement of >0.9mm or global BOLD signal changes >5 s.d. were flagged as potential outliers. Identified outliers were later included in the first-level statistical design. A new reference image, based on all scans except marked outliers is produced. The Structural and functional data are then realigned and normalised to MNI space, and segmented into grey matter, white matter and CSF, using SPM12 unified segmentation and normalisation (Ashburner & Friston, 2005). Default Conn parameters were used. Functional data were then smoothed using a Gaussian kernel of 8mm FWHM.

#### Condition vs Baseline Activation Maps

The functional imaging session was modelled at the single-subject level using a GLM in SPM12. The design consisted of one regressor per condition (Sentence Reading, Visual Commands, Sentence Listening, Auditory Commands, Horizontal Checkerboards, Vertical Checkerboards). For each participant, additional regressors were included to flag any outlier scans identified during realignment (one regressor per scan). Six regressors of no interest coding for scan-to-scan movement (x, y and z translations and rotations) as well as a seventh term coding for scan-to-scan global BOLD change, and a constant term were added. Stimulus blocks were modelled as epochs convolved with the canonical haemodynamic response function in SPM12, and rest trials (baseline) were left unmodelled. To rule out the possibility of systematic effects of participant movement on any literacy-related results, the number of identified outliers was tested for a relationship with word reading scores, no significant relationships were found (Time 1: Kendall’s ⊤ b = .086, p=.263; Time 2: ⊤ = -.094, p=.323). This also suggests that there was no systematic effect of literacy on compliance with the instruction to remain as still as possible in the scanner.

For each participant, parameter estimates for the auditory sentence and visual sentence conditions were contrasted with the baseline, and the resulting contrast images were used for second-level, random-effects, analyses, in which the individual subject data were tested for reliable group-level effects using a one-group t-test.

#### Relationship between BOLD Response and Literacy

Due to the non-normal distribution of the reading scores across the group, analyses that sought to probe a literacy - BOLD link were executed using non-parametric statistics. This was achieved using the Randomise tool from the FSL package (Winkler, Ridgway, Webster, Smith, & Nichols, 2014), using 5000 permutations to determine the null-distribution of the statistic for the contrast of interest. Results reported are significant at a cluster-mass threshold of p<.05 FDR-corrected for multiple comparisons, with a cluster-forming threshold set to be equivalent to uncorrected *p*<.001 (for N=91, t_(90)_=3.092)

#### Functional connectivity

Functional connectivity analyses were carried out using the CONN toolbox version 20.b (Whitfield-Gabrieli & Nieto-Castanon, 2012). First level design matrices for each participant from the fixed-effects analysis described above were entered into the toolbox. The default denoising pipeline was also run, with temporal filtering adjusted to use a high-pass filter with a cutoff of 0.008Hz. The low-frequency cut-off was omitted since the experimental conditions were presented in a block design, with relatively long block durations. The denoising procedure is fully described in Whitfield-Gabrieli and Nieto-Castanon (2012).

First-level functional connectivity was estimated for the three ROIs (PT, VWFA and pSTG) with the rest of the brain (seed-to-voxel connectivity). This involves the calculation of a Fisher-transformed bivariate correlation coefficient between the BOLD time-series of the ROI and the BOLD time-series of each of the non-ROI voxels. In order to probe the relationship between connectivity and literacy, the individual condition-wise connectivity maps were then tested, voxelwise, for a correlation with literacy, as indexed by word-reading scores. In order to avoid errors due to the non-normal distribution of the reading scores, permutation testing (5000 permutations), as implemented in CONN for cluster-mass thresholding, was used to determine significance of any effects at a whole-brain FDR corrected level of *p*<.05, with a cluster-forming threshold of uncorrected *p*<.001.

### Literacy Training Intervention and MRI Follow-Up

A number of the illiterate participants (N=22) completed a six-month literacy training program, after which they participated in an MRI session of the same design and with the same stimuli as described above. Twelve illiterate participants returned for a follow-up scan without training as did 25 literate participants. The training and non-training groups were matched for age, gender, handedness, income, number of literate family members, reading scores and non-verbal intelligence (Table 3). Complete details of the training procedure and its efficacy are described elsewhere (Hervais-Adelman et al., 2019), and summarised in the supplementary materials. The longitudinal analyses were confined to the evaluating the response of the ROIs to auditory and visual sentences.

**Table 3:** Characteristics of follow-up participants.

|  | **Trainees** | **No Train** | **Literate** |
| --- | --- | --- | --- |
| **N:** | 22 | 12 | 26 |
| *Age (Range)* | 31.36 (22-40) | 30.83 (22-49) | 26.84 (18-40) |
| *Sex [F:M]* | 21:1 | 10:2 | 8:17 |
| *Handedness [R:L]* | 22:0 | 12:0 | 25:0 |
| *Income (Range) [IRN/Month]* | 1795.455 (0-3000) | 2500 (2000-3000) | 1823.92 (0-9000) |
| *Marital Status*<br>*(Married:Unmarried:Widowed)* | 21:0:1 | 12:0:0 | 16:08:01 |
| *Number of Family Members* | 4.955 (2-8) | 4.667 (2-7) | 4.6 (2-8) |
| *Number of Literate Family Members* | 2.714 (0-6) | 2.889 (1-4) | 1.944 (1-5) |
| *Years of Schooling (Range)* | 0 (0) | 0 (0) | 4.88 (0-11) |
| *Word Reading Score T1*<br>*(Range) [maximum 86]* | 0.545 (0-6) | 3.25 (0-13) | 65.48 (9-86) |
| *Word Reading Score T2*<br>*(Range) [maximum 86]* | 7.591 (0-33) | 3.417 (0-14) | 68.44 (19-85) |
| *Letter recognition Score T1*<br>*(Range) [maximum 46]* | 9.227 (0-38) | 9.417 (0-35) | 39.04 (20-46) |
| *Letter recognition Score T2*<br>*(Range) [maximum 46]* | 33.227 (30-46) | 7.33 (0-36) | 41.76 (30-46) |
| *Raven's Matrices (Range)* | 13 (8-18) | 13.167 (8-27) | 17.8 (10-35) |

**Table 4:** Sentence listening vs Baseline. Loci of peaks (local maxima within a cluster separated by a minimum of 8mm) showing significant (cluster-size p<.05 FDR-corrected for multiple comparisons with a cluster-forming threshold of p<.001uncorrected) increase in BOLD over all participants when listening to sentences. Bold rows indicate peak of a cluster. Abbreviations: Hem-Hemisphere, L-Left; R-Right; unc. – uncorrected; fdr – false discovery rate

| Hem | Label | Brodmann Area | Cluster Size (voxels) | Cluster size $p(\text{unc.})$ | Cluster Size $p(\text{fdr})$ | Coordinates (MNI) | | | Statistic T | Voxel $p(\text{fdr})$ |
| --- | --- | --- | --- | --- | --- | --- | --- | --- | --- | --- |
|  |  |  |  |  |  | x | y | z |  |  |
| R | **Superior Temporal Gyrus** | 22 | 5533 | <.001 | <.001 | 60 | -10 | 2 | 23.55 | <.001 |
| R | Superior Temporal Gyrus | 22 |  |  |  | 58 | -18 | 4 | 22.85 | <.001 |
| R | Heschl's Gyrus |  |  |  |  | 48 | -18 | 10 | 21.36 | <.001 |
| L | **Superior Temporal Gyrus** | 22 | 6661 | <.001 | <.001 | -54 | -14 | 6 | 23.16 | <.001 |
| L | Superior Temporal Gyrus | 22 |  |  |  | -62 | -20 | 6 | 21.97 | <.001 |
| L | Superior Temporal Gyrus | 22 |  |  |  | -52 | -2 | -6 | 17.21 | <.001 |
| L | Superior Temporal Pole | 21 |  |  |  | -46 | 4 | -16 | 13.39 | <.001 |
| L | White Matter (splenium of Corpus Callosum) |  |  |  |  | -14 | -34 | 22 | 4.57 | .036 |
| R | **Cerebellum Lobule 8** |  | 339 | .001 | .005 | 24 | -62 | -52 | 8.63 | <.001 |
| R | Cerebellum Lobule 7 |  |  |  |  | 18 | -76 | -42 | 5.19 | .005 |
| L | **Inferior Frontal Gyrus Pars Triangularis** | 47 | 258 | .004 | .011 | -46 | 30 | 0 | 4.59 | .036 |
| L | Inferior Frontal Gyrus *Pars Orbitalis* | 47 |  |  |  | -40 | 34 | -12 | 4.2 | .09 |

**Table 5:** Sentence Reading vs Baseline. Loci of peaks (local maxima within a cluster separated by a minimum of 8mm) showing significant (cluster-size p<.05 FDR-corrected for multiple comparisons with a cluster-forming threshold of p<.001uncorrected) increase in BOLD for visual sentence presentation vs baseline over participants when listening to sentences. Bold rows indicate peak of a cluster. NB that these sentences did not constitute linguistically meaningful stimuli to the illiterate participants. Abbreviations: Hem-Hemisphere, L-Left; R-Right; unc. – uncorrected; fdr – false discovery rate

| Hem | Label | Brodmann Area | Cluster Size (voxels) | Cluster size $p$(unc.) | Cluster Size $p$(fdr) | Coordinates (MNI) | | | Statistic T | $p$(fdr) |
| --- | --- | --- | --- | --- | --- | --- | --- | --- | --- | --- |
|  |  |  |  |  |  | x | y | z |  |  |
| R | **Middle Occipital Gyrus** | **18** | **12509** | **<.001** | **<.001** | **28** | **-86** | **0** | **10.49** | **<.001** |
| L | Middle Occipital Gyrus | 18 |  |  |  | -24 | -90 | 6 | 10.16 | <.001 |
| L | Middle Occipital Gyrus | 18 |  |  |  | -28 | -86 | -2 | 9.9 | <.001 |
| L | Posterior Fusiform Gyrus | 18 |  |  |  | -20 | -86 | -6 | 9.9 | <.001 |
| L | Inferior Occipital Gyrus | 19 |  |  |  | -42 | -76 | -6 | 9.86 | <.001 |
| R | Superior | 18 |  |  |  | 22 | -90 | 10 | 9.52 | <.001 |
| R | Inferior Occipital Gyrus | 19 |  |  |  | 40 | -80 | 0 | 9.52 | <.001 |
| R | Fusiform Gyrus | 37 |  |  |  | 38 | -52 | -12 | 8.41 | <.001 |
| L | Posterior Fusiform Gyrus | 19 |  |  |  | -32 | -74 | -12 | 8.22 | <.001 |
| R | Inferior Occipital Gyrus | 37 |  |  |  | 36 | -66 | -8 | 8.14 | <.001 |
| L | Fusiform Gyrus | 37 |  |  |  | -42 | -56 | -10 | 7.86 | <.001 |
| L | Fusiform Gyrus | 37 |  |  |  | -36 | -42 | -18 | 6.64 | <.001 |
| L | Inferior Occipital Gyrus | 19 |  |  |  | -28 | -84 | 20 | 6.47 | <.001 |
| R | Fusiform Gyrus | 37 |  |  |  | 34 | -40 | -22 | 6.36 | <.001 |
| R | Hippocampus | 20 |  |  |  | 30 | -28 | -4 | 5.11 | 0.007 |
| R | Hippocampus | 20 |  |  |  | 34 | -12 | -20 | 4.57 | 0.036 |
| R | Middle Occipital Gyrus | 19 |  |  |  | 32 | -62 | 30 | 4.12 | 0.116 |
| R | Hippocampus | 20 |  |  |  | 36 | -22 | -14 | 3.75 | 0.322 |
| L | Fusiform Gyrus | 30 |  |  |  | -24 | -32 | -16 | 3.69 | 0.364 |
| L | Inferior Parietal Cortex | 7 | 334 | 0.001 | 0.006 | -24 | -48 | 48 | 5.87 | 0.001 |
| R | Inferior Parietal Cortex | 7 | 220 | 0.006 | 0.021 | 28 | -48 | 54 | 5.72 | 0.001 |

**Table 6:**
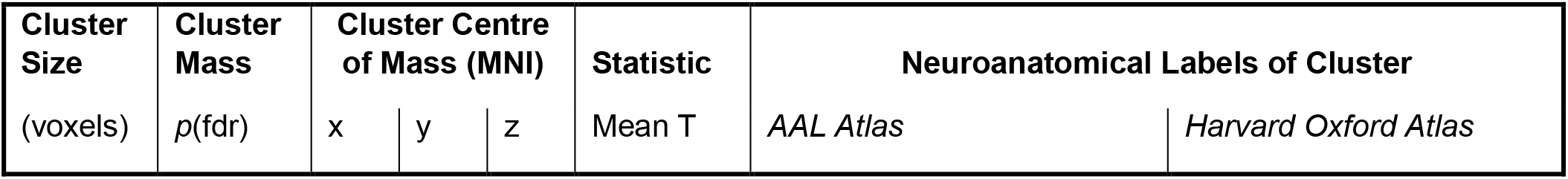
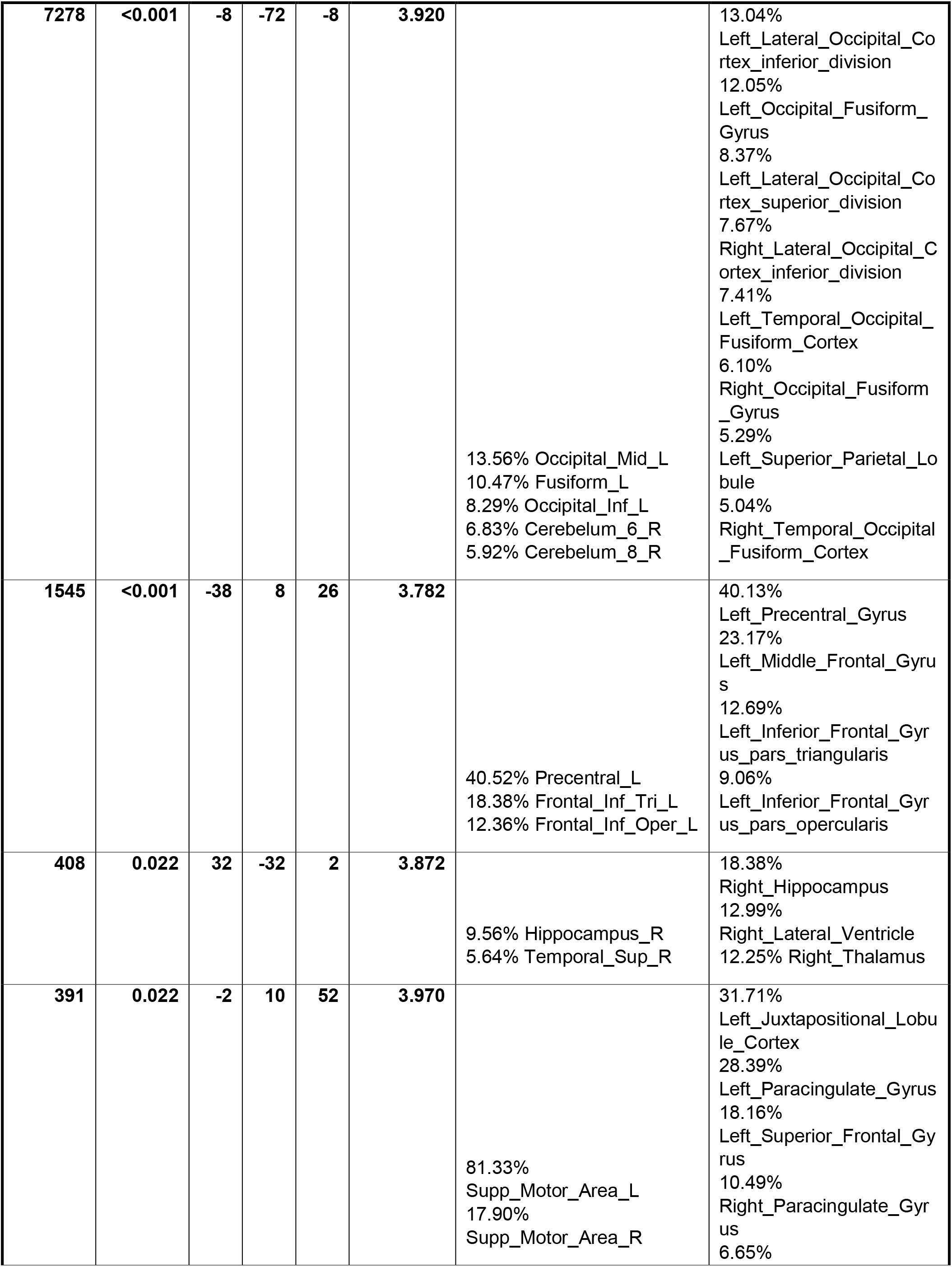
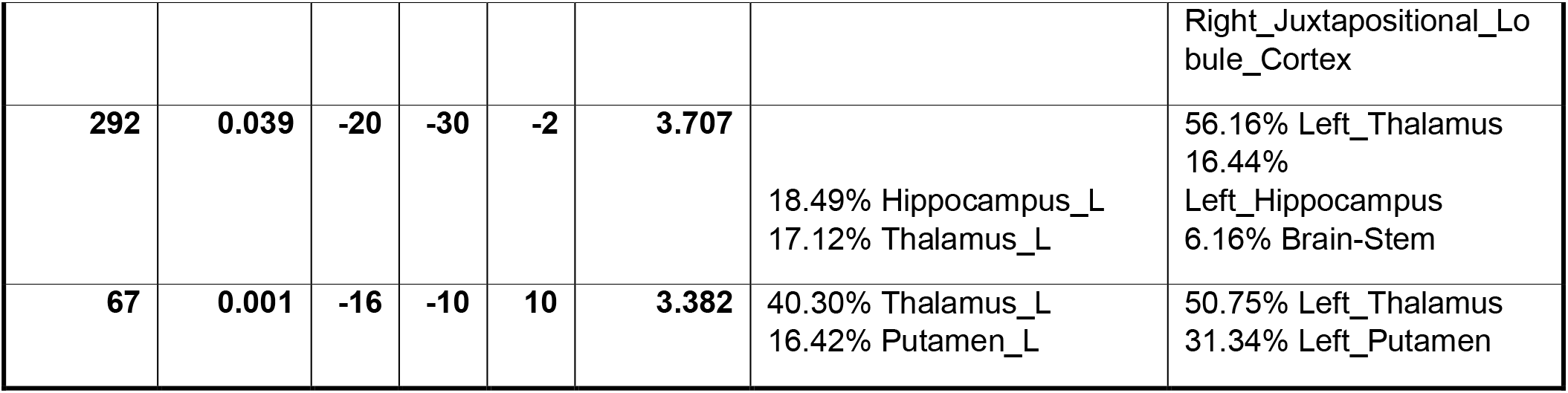
Modulation of Responses to Written Sentences by Literacy: Loci of positive associations between brain response to visually-presented sentences and literacy, assessed by word-reading scores. Cluster-Mass p<.05 FDR-corrected for multiple comparisons with a cluster-forming threshold of p<.001 uncorrected. Results are tabulated by cluster and the proportion of coverage of neuroanatomical regions are provided for two atlases (see methods for details). Coverages do not always sum to 100%, as atlases do not label all areas in the brain volume (e.g. white matter, some subcortical areas, CSF, ventricles)

**Table 7:** Significant modulation of pSTG connectivity by literacy during auditory sentence presentation. Cluster-Mass p<.05 FDR-corrected for multiple comparisons with a cluster-forming threshold of p<.001 uncorrected. Results are tabulated by cluster and the proportion of coverage of neuroanatomical regions are provided for two atlases (see methods for details). Coverages do not always sum to 100%, as atlases do not label all areas in the brain

| Cluster Size<br>(voxels) | Cluster Mass<br>$p(\text{fdr})$ | Cluster Centre of Mass (MNI) | | | Statistic<br>Mean T | Neuroanatomical Labels of Cluster | |
| --- | --- | --- | --- | --- | --- | --- | --- |
|  |  | x | y | z |  | AAL Atlas | Harvard Oxford Atlas |
| 602 | .046 | -24 | -14 | 58 | 3.564 | 80.33%<br>Precentral_L<br>10.83%<br>Frontal_Sup_2_L<br>5.33%<br>Postcentral_L | 74.67% Left_Precentral_Gyrus<br>16.50% Left_Postcentral_Gyrus<br>8.83%<br>Left_Superior_Frontal_Gyrus |

**Table 8.** Significant modulation of VWFA connectivity by literacy during visual sentence presentation. Loci of positive associations between literacy and effective connectivity of VWFA during visual sentence presentation. Cluster-level p<.05 FWE-corrected for multiple comparisons with a cluster-forming threshold of p<.001 uncorrected. Bold rows indicate peak of a cluster. Abbreviations: Brodmann Area; Hem-Hemisphere, L-Left; R-Right; FWE –Familywise Error

| Cluster Size<br>(voxels) | Cluster Mass<br>$p(\text{fdr})$ | Cluster Centre of Mass (MNI) | | | Statistic<br>Mean T | Neuroanatomical Labels of Cluster | |
| --- | --- | --- | --- | --- | --- | --- | --- |
|  |  | x | y | z |  | AAL Atlas | Harvard Oxford Atlas |
| 1074 | <.001 | -48 | 26 | 0 | 4.244 | 61.33%<br>Frontal_Inf_Tri_L<br>26.03%<br>Frontal_Inf_Orb_2_L<br>9.93%<br>Frontal_Inf_Oper_L | 40.36%<br>Left_Inferior_Frontal_Gyrus_<br>pars_triangularis<br>27.90%<br>Left_Inferior_Frontal_Gyrus_<br>pars_opercularis<br>20.60%<br>Left_Frontal_Orbital_Cortex<br>10.21%<br>Left_Frontal_Operculum_Cor<br>tex |
| 540 | <.001 | -46 | 0 | 46 | 4.138 | 89.07% Precentral_L<br>6.48% Frontal_Mid_2_L | 65.37%<br>Left_Precentral_Gyrus<br>34.63%<br>Left_Middle_Frontal_Gyrus |
| 507 | 0.010 | -18 | -90 | -4 | 3.403 | 55.34% Occipital_Mid_L<br>21.54% Occipital_Inf_L<br>8.50% Fusiform_L<br>6.72% Lingual_L | 41.50% Left_Occipital_Pole<br>34.58%<br>Left_Occipital_Fusiform_Gyr<br>us<br>19.57%<br>Left_Lateral_Occipital_Corte<br>x_inferior_division |
| 493 | <.001 | -6 | 12 | 54 | 4.087 | 91.08%<br>Supp_Motor_Area_L<br>5.27%<br>Supp_Motor_Area_R | 55.58%<br>Left_Superior_Frontal_Gyrus<br>19.27%<br>Left_Juxtapositional_Lobule_<br>Cortex<br>19.07%<br>Left_Paracingulate_Gyrus |
| 267 | 0.016 | 38 | -62 | -26 | 3.931 | 49.81% Cerebelum_6_R<br>40.82%<br>Cerebelum_Crus1_R<br>9.36% Fusiform_R | 24.34%<br>Right_Occipital_Fusiform_Gy<br>rus<br>16.10%<br>Right_Temporal_Occipital_F<br>usiform_Cortex |
| 189 | 0.041 | 38 | -84 | 12 | 3.498 | 95.08% Occipital_Mid_R | 38.80%<br>Right_Lateral_Occipital_Cort<br>ex_inferior_division<br>38.80%<br>Right_Lateral_Occipital_Cort<br>ex_superior_division; |
| 189 | 0.037 | -56 | -34 | 4 | 3.723 | 87.23%<br>Temporal_Mid_L<br>9.04% no_label | 82.45%<br>Left_Superior_Temporal_Gyr<br>us_posterior_division<br>9.04%<br>Left_Middle_Temporal_Gyru<br>s_posterior_division<br>5.85%<br>Left_Planum_Temporale |

#### Pre- vs Post-Training Comparison of pSTG Connectivity

The connectivity analysis was re-run for all participants who returned for a second fMRI scan (N=60). The connectivity values, estimated as Beta weights output by the CONN toolbox, were extracted for a representative sphere of radius 8mm centred at the centre of mass of the GFMA cluster showing literacy-modulated connectivity with pSTG during speech processing (illustrated in Figure 3, see also Table S7). The mean connectivity within this sphere was calculated as the mean of all voxels (ignoring any missing values). The mean values for Timepoint 1 and Timepoint 2 were then compared using a repeated measures ANVOA with a between-participants factor of group and a within-participants factor of timepoint. Post-hoc pairwise comparisons were executed and corrected using Holm’s method for the full family (N=15) of comparisons executed.

#### Additional Bayesian Analyses of BOLD-Literacy Correlations in ROIs

Given the partially confirmatory nature of this investigation (attempting to reproduce effects of literacy on the response of auditory phoneme-processing cortical regions during speech processing), where possible, Bayesian analyses were executed to estimate the strength of any failures to replicate the prior effect. Bayesian analyses were carried out using JASP (JASP Team), using default parameters, unless otherwise indicated. Frequentist statistics are reported for the whole-brain fMRI analyses, as Bayesian analysis tools are not yet readily available for these. Frequentist p values are provided wherever Bayes factors are not available. Bayes factors are presented as BF_10_, indicating the ratio of evidence in favour of the alternative hypothesis. Values greater than one indicate evidence in favour of the alternative hypothesis, values lower than one indicate evidence in favour of the null hypothesis.

## Results

### Relationships between reading ability and demographic factors

Complete demographic data are presented in Supplementary Table 1. These data are also presented in a previous publication reporting on these participants *(38)*.

Pairwise correlation analyses using Kendall’s ⊤ b were carried out to test for relationships between literacy and various demographic factors, including age, sex, monthly income and performance on Raven’s Matrices of all the participants at the first time point (N participants = 91). The complete set of pairwise correlations is reported in Supplementary Table 2. As would be expected from the participant selection procedure, significant correlations of interest were found between the literacy measures (word reading and Akshara recognition) and Years of schooling (Word reading: ⊤=.605, *p*<.001, Akshara recognition: ⊤=0.580, *p*<.001). There was also a significant relationship between literacy and sex (dummy coded as a binary variable), such that female participants were less likely to be literate, which is due to cultural factors affecting access to schooling at the study site (Word reading: ⊤=.420, *p*<.001, Akshara recognition: ⊤=0.411, *p*<.001). There was a negative correlation between age and literacy (Word reading: ⊤=-.214, *p*=.006 Akshara recognition: ⊤=-.228, *p*=.003), indicating that the more literate participants tended to be younger than the illiterate participants. A further relationship was found between literacy and Raven’s matrices performance (Word reading: ⊤=.242, *p*=.002, Akshara recognition: ⊤=0.278, *p*<.001).

In order to rule out a potential confounding effect of underlying fluid intelligence differences, a mediation analysis was carried out to test for a potential mediating effect of Raven’s Matrices performance on the relationship between years of schooling and literacy. The analysis was carried out using the MeMoBootR package *(Buchanan, 2018)* in R version 4.0.5 *(R Core Team, 2015)*. The path plot for the mediation analysis is shown in Figure 1. Evaluating the relationship between years of schooling mediated by Raven’s (assumed to be a proxy for non-verbal IQ) shows a direct effect of schooling on word reading (b = 6.76, t(89)=9.362, p<.001), a significant relationship between Education and Raven’s (b = 0.402, t(89)=2.74, p=.007) and a significant relationship between Raven’s and Literacy when accounting for Education (b = 1.182, t(88)=2.319, p=.023). The direct effect of Education on Literacy is, however, not significantly modulated by Raven’s (direct effect, accounting for Ravens, b = 6.285, t(88) = 8.562, p<.001, Aroian Sobel test z=1.705, p=.088, bootstrapped 95% confidence for the indirect effect: −0.151, 1.130). Thus, the component of non-verbal intelligence reflected in Raven’s performance is not responsible for literacy and it may even be speculated that superior Raven’s performance in the literate participants could itself be the result of schooling.

**Figure 1.**
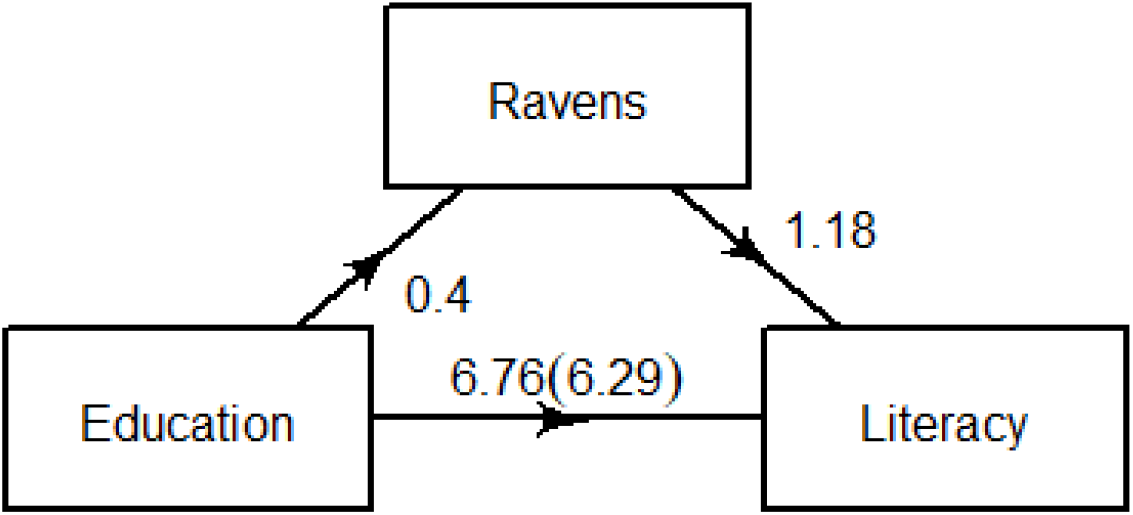
Path plot for mediation analysis of effect of Raven’s performance on the direct relationship between years of schooling and literacy. Mediation analysis suggests that the influence of years of schooling on literacy is not mediated by Raven’s performance (Sobel test z=1.705, p=.088, bootstrap (N=1000 permutations) 95% confidence interval: −0.151, 1.130)

It is noteworthy that monthly income is not significantly related to literacy (Word reading: ⊤=.119, *p*=.147, Akshara recognition: ⊤=.108, *p*<.171), indicating that literacy is unlikely to be a primary determinant of socio-economic status in the communities from which the study participants were drawn.

### Six months of Literacy Training – A partially successful intervention

Of the 91 participants initially tested, 60 returned for a follow-up scan, for whom complete data are available for only 59 (due to an error in collecting behavioural data for one participant during the follow-up session), comprising three groups: Literate (N=25, mean Word reading score: 64.58), Illiterate participants who participated a six-month literacy training and non-trained (N=22, mean word reading score: 0.55), control illiterate individuals (N=12, mean word reading score: 3.25).

Repeated measures ANOVA, including a between-subjects factor of Group (Literate, Illiterate Trainee and Illiterate Control) and a within-participants factor of Session (Time 1, Time 2) was used to determine whether training had a significant impact on literacy. As reported previously (Hervais-Adelman et al., 2019), the literacy training program was a mixed success, descriptive plots are shown in Figure 2. A significant group-by-timepoint interaction indicated that trained participants improved in their Akshara recognition performance (F_(2,56)_=58.463, *p*<.001,partial-η^2^=.676), but this interaction was not significant for word reading (F_(2,56)_=1.430, *p*=248, partial-η^2^= .049). Planned pairwise tests for the simple effect of Time on literacy scores within group indicated significant improvements in the trainees (*N*=22, Akshara: F_(1)_=200.69, *p*<.001; Word reading: F_(1)_=7.673, *p*=.008) that were absent in the untrained illiterate participants (*N*=12, Akshara: F_(1)_=0.825, *p*=.368; Word reading: F_(1)_=0.002, *p*=.962) and in the Literate participants, who also received no training (*N*=25, Akshara: F_(1)_=2.929, *p*=.093; Word reading: F_(1)_=1.539, *p*=.220). These results confirm that training improved literacy, albeit that the six-month program did not suffice to render the formerly illiterate individuals fluent readers of words in Devanagri script.

**Figure 2.**
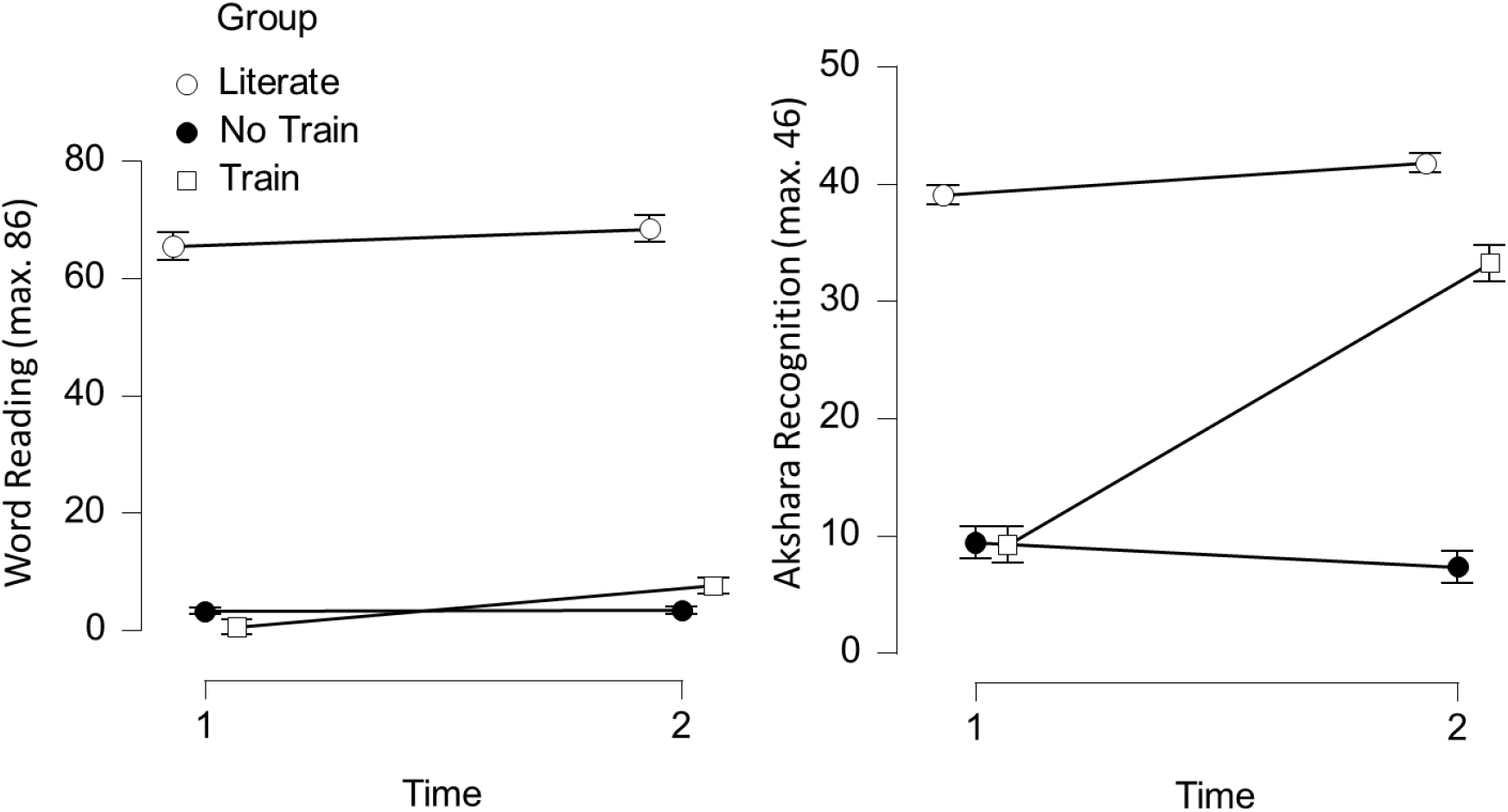
Mean reading scores by participant group, at Times 1 and 2. Error bars show standard error of the mean. Left panel shows word reading performance, right panel shows Akshara recognition performance. A significant improvement in Akshara recognition, and to a lesser extent, word reading performance, is found in the trained illiterate participants.

### Cerebral Responses to Speech and Orthographic Stimuli

Initial control analyses comparing the sentence listening and sentence reading conditions to baseline served to verify that expected patterns of, respectively, auditory, and visual activation were apparent (Figure 3). Listening to sentences produced significant increases in substantial expanses of bilateral superior temporal areas, consistent with speech processing (Hickok, 2012; Price, 2012), as well as increases in left inferior frontal gyrus broadly consistent with linguistic processes implicated in sentence processing (Friederici, 2012; Hagoort, 2017; Price, 2012). Visual presentation of sentences (which, it must be noted, were not interpretable for a large proportion of the participants) lead on average to widespread significant bilateral visual cortical activation compared to baseline.

**Figure 3.**
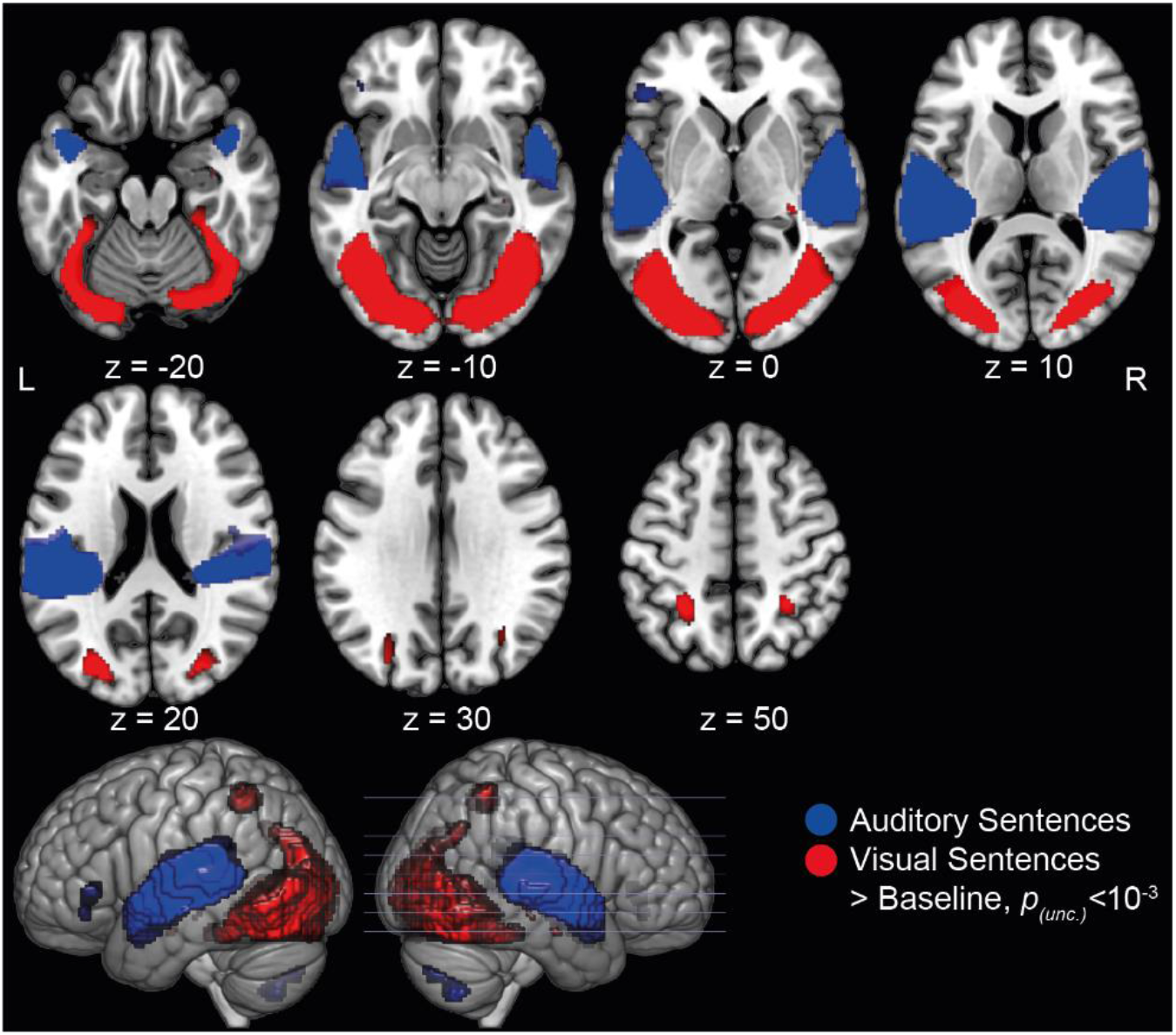
Group mean activation map for Auditory Sentences vs Baseline and Visual Sentences vs Baseline, thresholded at voxelwise p(unc.)<.001, with a cluster-level significance of p(FWE)<.05, projected on MNI single subject template brain. Z co-ordinates (MNI mm) are supplied for each slice and marked on the render in bottom-row. Abbreviations: L – left, R - right

In order to determine whether literacy in Devanagari (functionally approximated by word reading score) has an impact on the BOLD response to speech, as previously reported for literacy in the Latin alphabet (Dehaene et al., 2010), we conducted a regression analysis to test for a correlation between word reading score and brain response during auditory sentence presentation and during visual sentence presentation.

#### No Evidence that Cerebral Responses to Speech are affected by Literacy

Testing for the effect of literacy on the patterns of BOLD activation revealed no loci at which there were significant relationships between word reading scores and brain response during sentence listening, at the relatively liberal threshold of uncorrected voxelwise *p*<.001.

Despite the absence of significant effects of literacy on the BOLD response to auditory sentences at whole-brain levels, it is conceivable that a more subtle relationship was missed. Given the existence of an *a priori* rationale for testing a specific ROI, we further probed the possibility that PT response to spoken sentences is modulated by literacy. We carried out a region of interest analysis in which the subjectwise mean PT response was extracted and tested for a relationship with literacy.

Bayesian correlation (Kendall’s ⊤ b) analysis was carried out to test the relationship between BOLD response to auditory sentence presentation in the two acoustic-phonetic ROIs (PT and pSTG), and literacy, quantified by Akshara recognition and word reading scores. These analyses revealed that there is no evidence in favour of relationships between literacy and brain response (PT, relationship with Akshara recognition: ⊤ = .095, BF_10_ = 0.332, relationship with Word reading, ⊤ = .007, BF_10_ = 0.137; pSTG relationship with Akshara recognition: ⊤ = -.090, BF_10_ = 0.303, relationship with Word reading: ⊤ = -.030, BF_10_ = 0.149). The associated Bayes factors indicate anecdotal to substantial levels of evidence in favour of the null hypothesis of no relationship between BOLD response and literacy. The ROIs and the relationships between BOLD response and word reading scores at the first and second time points are shown in Figure 4.

**Figure 4:**
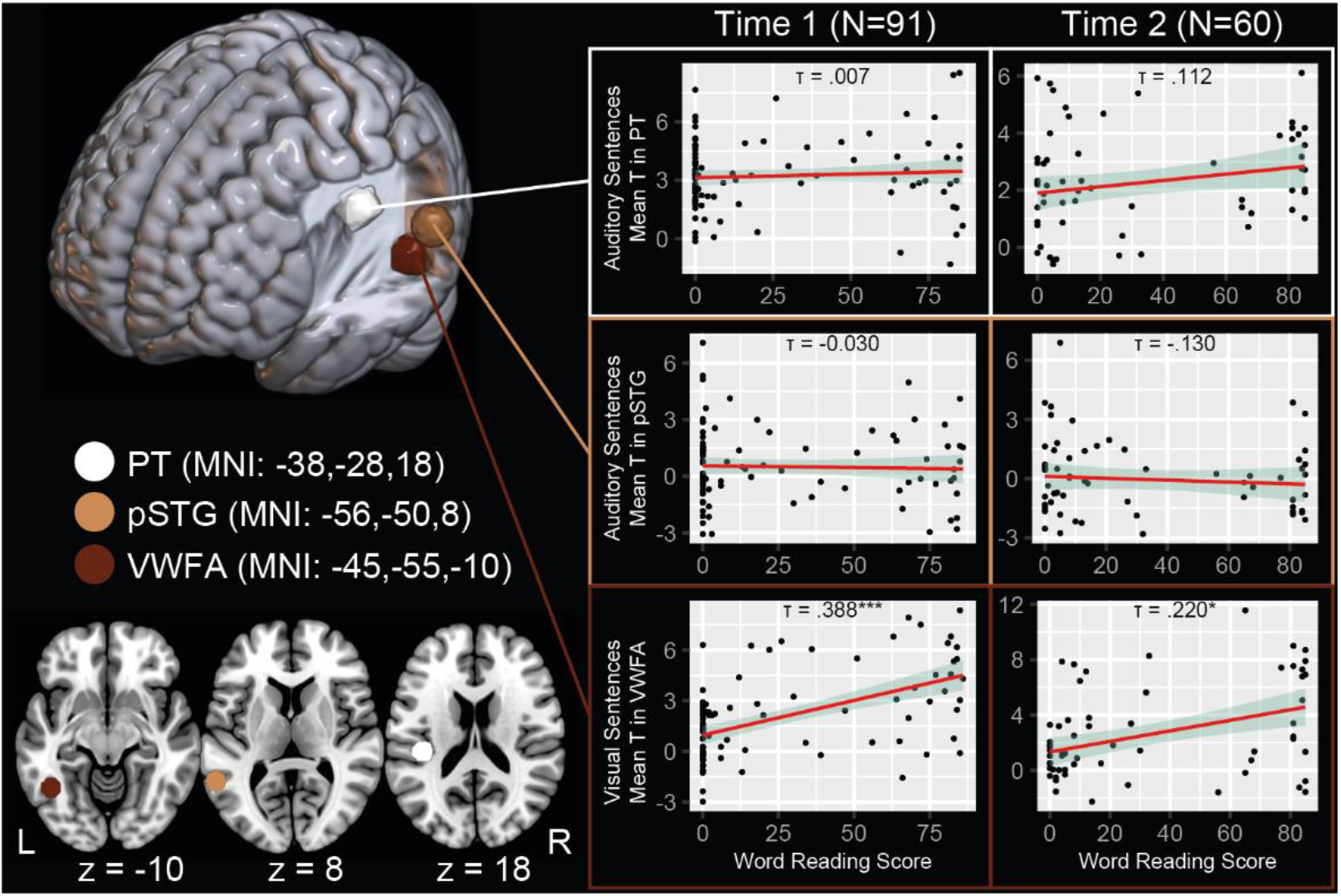
Relationship between word-reading scores and BOLD response to Auditory Sentences or Visual sentences, plotted for each of the three ROIs examined in this study (Planum Temporale, PT; Posterior Superior Temporal Gyrus, pSTG; Visual Wordform Area, VWFA). Scatter plots show individual subject mean T statistics in the sampled region at each timepoint of the study, trend line indicates fit of robust linear regression, ribbon indicates 95% CI of fit. Due to the non-normal distribution of the data for word reading, statistical analyses were carried out using non-parametric methods (Kendall’s ⊤ b), trendlines are illustrative. ⊤ values indicated on each scatter plot indicate Kendall’s ⊤ and statistical significance uncorrected for multiple testing *<.05, ***<.001. It is clear that there is a significant positive relationship between word-reading ability and response to visual sentences in VWFA, while there is no significant relationship between responses to auditory sentences and literacy in the two superior temporal ROIs.

#### Cerebral Responses to Orthographic Stimuli are Modulated by Literacy

As previously reported on these data (Hervais-Adelman et al., 2019), BOLD responses to visual sentence presentation was significantly modulated by literacy in a number of areas consistent with the reading network, implicating visual, occipito-fusiform, midline motor and left inferior frontal gyral regions (Figure 5).

**Figure 5:**
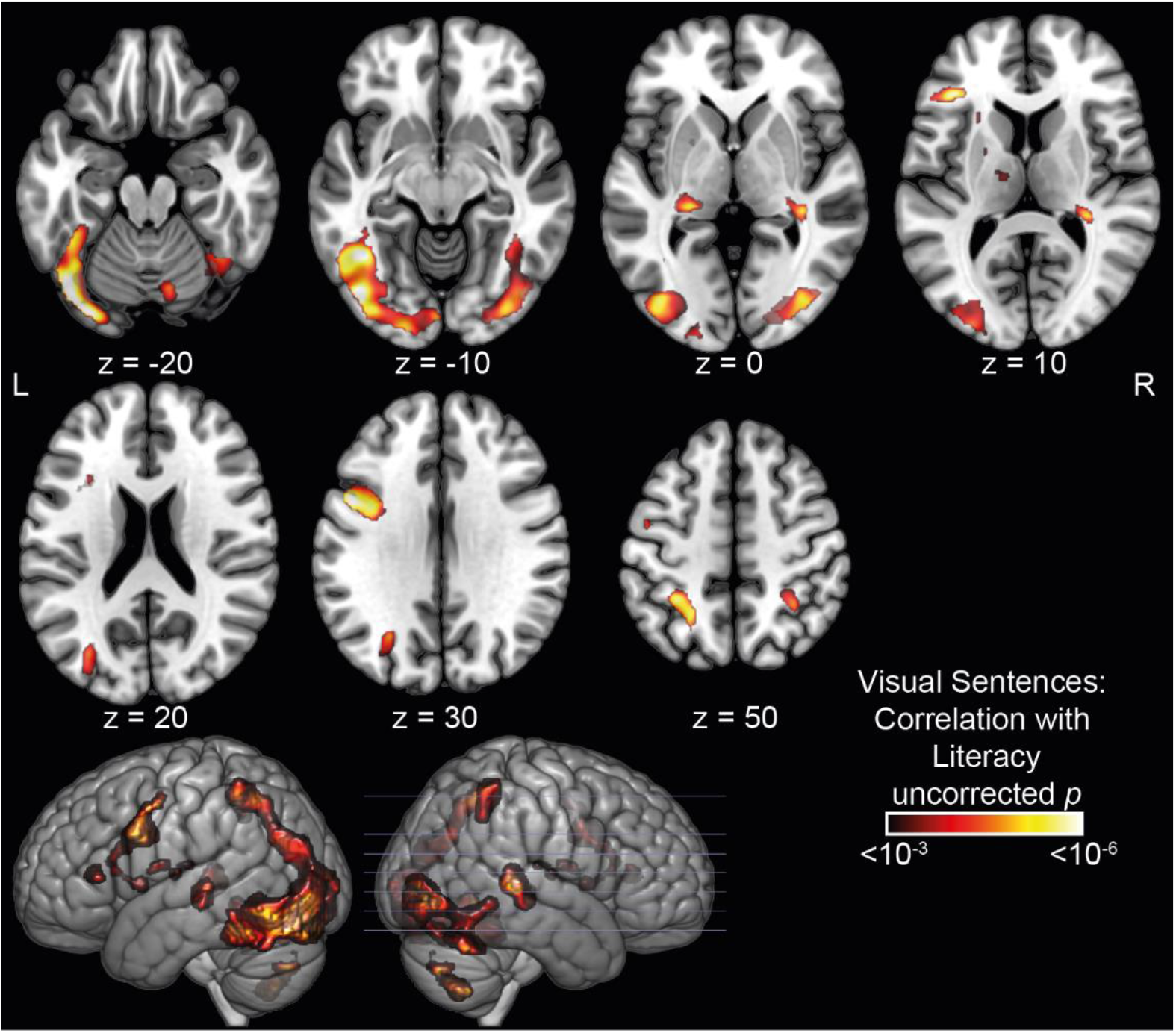
Correlation between BOLD response to Visual Sentences and literacy. (indexed by word reading score) at Time 1, thresholded using at cluster-mass p_(FDR)_<.05 with a cluster-forming threshold of p<.001. Results projected onto MNI single subject brain. Z co-ordinates (MNI mm) are supplied for each slice and marked on the render in bottom-row. Abbreviations: L – left, R - right

To ensure that the expected effect of literacy on brain response could be discerned for the BOLD response to orthographic stimuli, an analysis of the modulation of response in VWFA by literacy was carried out. A Bayesian correlation shows decisive evidence in favour of the existence of a correlation between literacy and BOLD response while reading sentences (Akshara recognition: Kendall’s ⊤= .354, BF_10_ = 27550.521; Word Reading: Kendall’s ⊤ = .388, BF_10_= 320744.068).

After training, participants showed significant improvements in Akshara recognition and marginally significant improvements in word reading performance (see Supplementary materials). The change in brain response to orthographic input as a function of literacy training in these participants has been described previously (Hervais-Adelman et al., 2019). Here we focus on three ROIs (PT, pSTG and VWFA) to establish whether within-participant improvements in reading ability affect the magnitude of the BOLD response to spoken and orthographic input.

#### Impact of Acquiring Literacy on Responses to Auditory and Visual Sentences in PT, pSTG and VWFA

An initial repeated measures ANOVA, with a within-participant factor of Session (Time 1, Time 2) and a between-participants factor of Group (Literate Control, Non-Trained Control, Trainee) was carried out to test for a significant group by time point interaction, which would be indicative of an effect of training. For completeness, this analysis was carried out on responses to both auditory and visual sentence presentation in all three ROIs.

In PT, no significant group-by-time point interaction was found for either listening to sentences (F_(2,57)_=0.590, p=.558, partial η^2^ = .020) or visually-presented sentences (F_(2,57)_=0.273, p=.762, partial η^2^ = .009), suggesting that there was no effect of training on BOLD response to auditory or orthographic sentences in this ROI. Similarly, there was no significant interaction between time and group for the pSTG ROI (auditory sentences: F(2,57)=0.596, p=.555, partial η^2^ = .020; visual sentences: F(2,57)=0.647, p=.527, partial η^2^ = .022). In VWFA, however, while there was no significant group-by-time point interaction for auditory sentences (F_(2,57)_=1.043, p=.359, partial η^2^ = .035), a significant group-by-time point interaction was found for visually presented sentences (F_(2,57)_=3.354, p=.042, partial η^2^ = .105), suggesting that literacy training selectively increased response to orthographic stimuli in the VWFA (This effect is illustrated in Figure 6).

**Figure 6:**
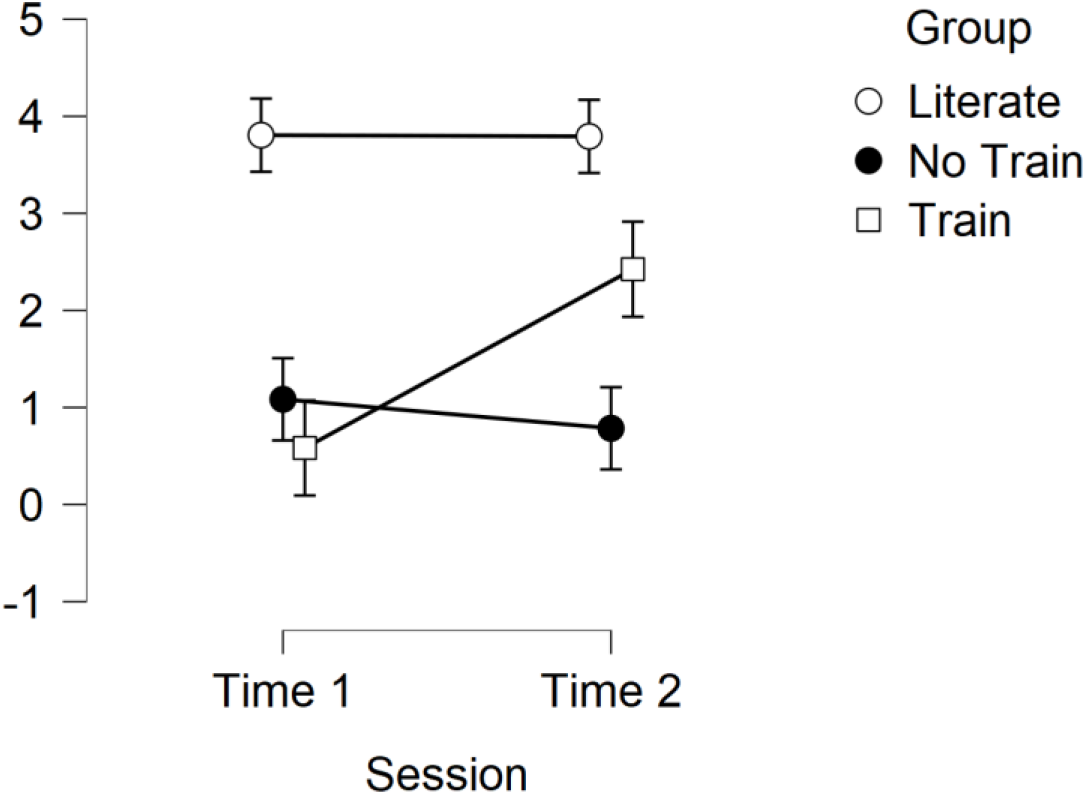
Mean response to Visual Sentences in VWFA ROI, by group and Session. Error bars show standard error of the mean. A literacy-training specific increase in responsiveness can be seen for the Trainee participants who received six months of literacy instruction between sessions.

A follow-up analysis (Bayesian paired t-test) was carried out on the null findings to determine the relative certainty that there was no categorical difference in brain response to the auditory or visual sentence materials as a result of training. This analysis was carried out for the PT and pSTG ROIs, comparing pre-training with post-training BOLD response to each condition and testing the directional hypothesis of post-training BOLD icnrease. These analyses revealed varying degrees of evidence in favour of the null hypothesis that there is **no** increase in response after training in either PT (auditory: BF_-0_= 0.056; visual: BF_-0_= 0.243), pSTG (auditory: BF_-0_= 0.076; visual: BF_-0_= 0.978). The same analyses in VWFA showed strong evidence in favour of the alternative hypothesis that BOLD response to visual sentences was greater after training (BF_-0_= 7.161), while there was substantial evidence in favour of the null hypothesis of no difference for the response to auditory sentences (BF_-0_= 0.311).

In sum, the analysis of within-subject change after training provides no evidence for an increase in recruitment of acoustic-phonetic related brain areas, while increasing literacy does lead to an increase in VWFA responsiveness to visually presented sentences.

#### Functional Connectivity of PT, pSTG and VWFA

It is self-evident that in hearing individuals the acquisition of literacy at some stage corresponds to acquiring a mapping of spoken language to orthographic symbols. This presumably implicates auditory processes and may therefore have functional consequences for regions associated with acoustic-phonetic processing. We therefore sought evidence for an impact of literacy on the functional connectivity of these regions, during auditory sentence presentation.

### Functional connectivity of PT during spoken sentence presentation is not modulated by literacy

Functional coupling to a seed region (the PT or pSTG ROI), during auditory sentence presentation was estimated and tested for modulation by literacy. We found no indication that there was a significant impact of literacy (even at a liberal voxelwise threshold of uncorrected *p*<.001) on the functional connectivity of the PT ROI.

### Functional Connectivity between pSTG and graphemic/motor frontal area during auditory sentence presentation increases as a function of literacy

A substantial cluster of voxels in dorsal frontal cortex (spanning Brodmann areas 6d, 4a and 4p) showed functional coupling to pSTG during sentence listening that was significantly positively associated with literacy (Figure 7). This cluster of dorsal sensorimotor voxels intersects with an area commonly said to have been identified by Exner (Exner, 1881; Roux, Draper, Köpke, & Démonet, 2010) as crucial for handwriting and more recently, repeatedly associated with handwriting in a number of functional imaging (Longcamp, Anton, Roth, & Velay, 2003; Longcamp et al., 2008; Longcamp et al., 2014; Planton, Longcamp, Péran, Démonet, & Jucla, 2017) and cortical stimulation studies (Roux et al., 2009). This result suggests that while learning to read may not necessarily alter the responsiveness of planum temporale to auditory sentences, a functional relationship between the pSTG acoustic-phonetic processing area and handwriting representations develops. The important implication is that learning to associate Devanagari characters with their acoustic-phonetic form appears to lead to the development of an auditory to graphomotor mapping.

**Figure 7:**
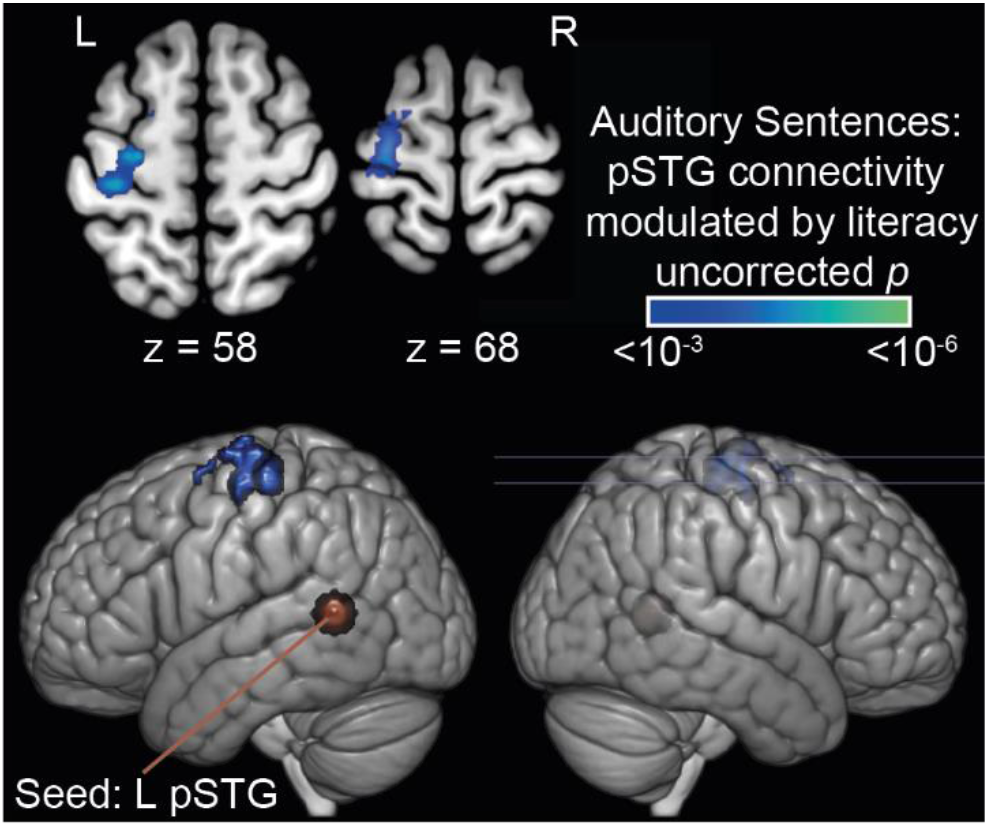
Significant correlation between functional connectivity of pSTG and literacy during auditory sentence processing. Results thresholded at cluster-mass p(FDR)<.05, using a cluster-forming threshold of voxelwise p<.001(single-tailed distribution, considering positive relationships only). Results projected onto MNI single subject brain. Z co-ordinates (MNI mm) are supplied for each slice and marked on the render in bottom-row. Abbreviations: L – left, R - right

### Functional connectivity of VWFA to auditory and prefrontal cortices during visual sentence presentation is modulated by literacy

As an additional control, the connectivity of VWFA was examined during sentence reading and the correlation between connectivity and literacy was evaluated. This revealed a broad pattern of connectivity between VWFA and the wider reading network (Figure 8), including the left inferior frontal gryus, left dorsal premotor cortices and supplementary motor areas, alongside bilateral visual areas and a region of posterior superior temporal sulcus associated with processing of speech. This latter cluster is somewhat distant, both neuroanatomically and in terms of auditory-processing (Rutten, Santoro, Hervais-Adelman, Formisano, & Golestani, 2019) from the planum temporale ROI examined in this study, but abuts the pSTG ROI defined *a priori*.

**Figure 8:**
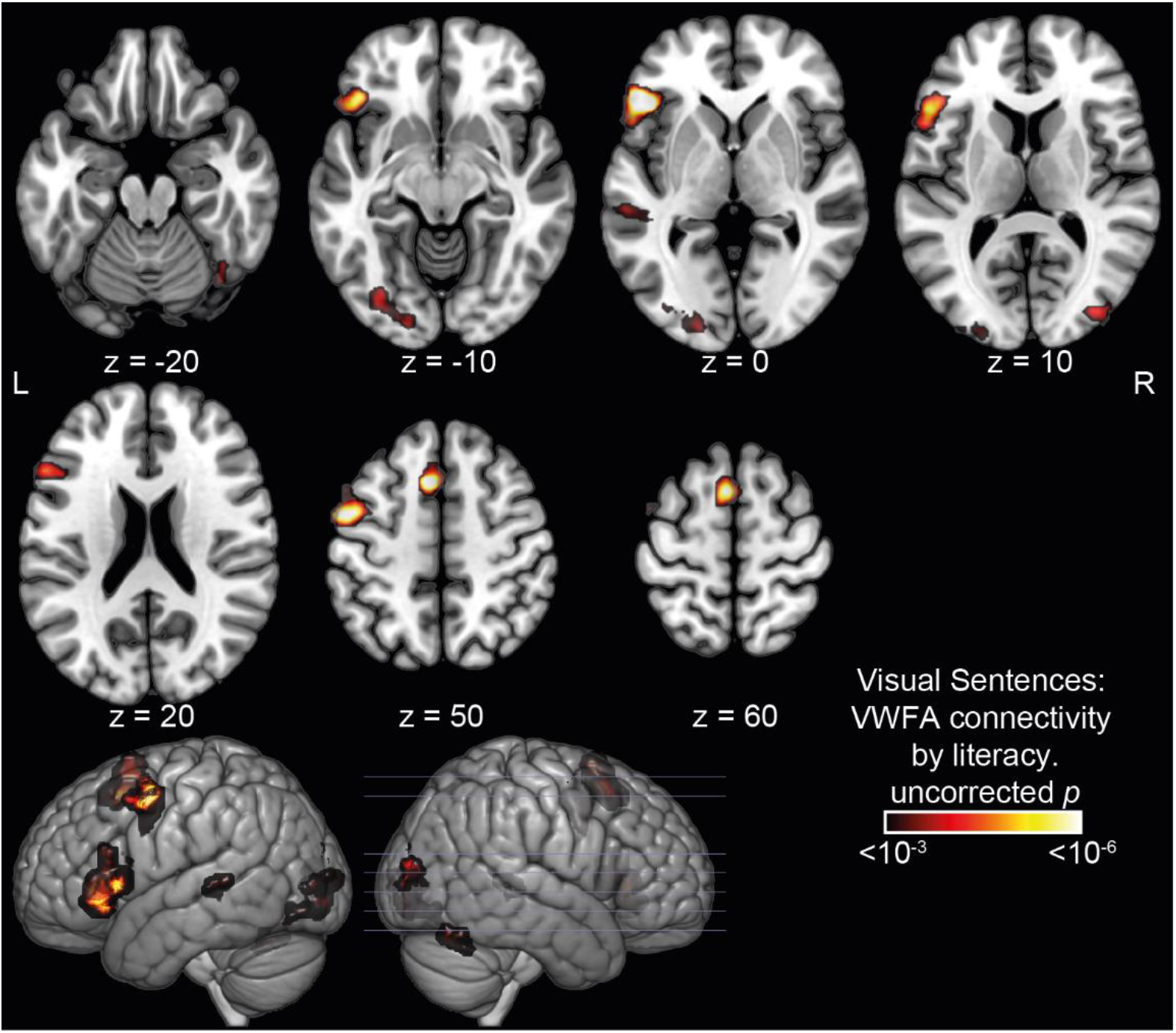
Brain areas showing significant modulation of VWFA connectivity by literacy during visual sentence processing. Results thresholded at cluster-mass p(FDR)<.05, using a cluster-forming threshold of voxelwise p<.001 (single-tailed distribution, considering positive relationships only). Results projected onto MNI single subject brain. Z co-ordinates (MNI mm) are supplied for each slice and marked on the render in bottom-row. Abbreviations: L – left, R - right

#### Literacy training increases functional connectivity of pSTG and GMFA during auditory sentence presentation

If learning to read is responsible for the observed correlation between literacy and functional connectivity of pSTG with GMFA during auditory sentence presentation, then the literacy training intervention should drive an increase in this connectivity. Connectivity estimates for all participants who returned for a second scan were estimated for the pSTG. Individual connectivity parameters at the centre of mass of the cluster identified as GMFA above (using a sphere of radius 8mm centred at MNI: −14, −24, 58) were extracted and tested for an effect of training using a repeated measures ANOVA (within-participants factor: Time 1, Time 2; between-participants factor: Literate Control, Untrained Control, Trainee). This analysis revealed a significant group-by-time comparison (F_(2,57)_=4.602, *p*=.015, partial η^2^ = .139), with post-hoc pairwise comparison showing that the effect is driven by an increase in connectivity in the trainees (t=3.324, *p*_holm_=.023), which was not found in the other groups (Literate: t=-0.331; Untrained: t=-.607, both *p*_holm_>.999).

## Discussion

Surprisingly, given the dominant assumption that literacy necessarily induces changes to online speech processing, we find no evidence for changes to online speech processing, either cross-sectionally or longitudinally, with Bayesian analyses suggesting substantial evidence in favour of there being no effect. However, there is compelling evidence that a functional connection during speech processing arises between pSTG and graphomotor areas, as a result of literacy.

These findings contrast starkly with those previously presented for literate and ex-illiterate readers of alphabetic script (Dehaene et al., 2010), who showed modulation of PT response during sentence listening as a function of literacy status. It was hypothesised that this is due to enhanced phonological processing, which is engaged in an obligatory manner during continuous speech processing. We find no such evidence, calling into question the *generalisability* of the hypothesis of radical reconfiguration of the functional role of auditory processing areas by the acquisition of literacy. Not only is there no whole brain level nor ROI-level effect of literacy on brain response to auditory sentence presentation, there is substantial evidence (at the ROI level) in favour of the hypothesis that there is no impact of literacy.

Nevertheless, we do not seek to call into question the substantial body of evidence that points towards behavioural consequences of literacy for speech processing tasks. However, the limited evidence that orthographic knowledge *can* affect the processing of spoken words comes from metalinguistic tasks that are completely or mostly offline such as rhyme judgments (Seidenberg & Tanenhaus, 1979), phoneme monitoring (Dijkstra, Roelofs, & Fieuws, 1995; Halle, Chereau, & Segui, 2000) and word blending (Ventura, Kolinsky, Brito-Mendes, & Morais, 2001), which require participants explicitly to breakdown (individual) spoken words into smaller units. Some other behavioural studies finding evidence for orthography-on-speech effects have used tasks that do not require meta-phonological judgments but which are clearly meta-linguistic in nature, such as auditory lexical decision and shadowing (Chereau, Gaskell, & Dumay, 2007; Ventura, Morais, Pattamadilok, & Kolinsky, 2004) and are far removed from how people ordinarily listen to spoken language. The few behavioural studies that have used online speech tasks suggest that orthographic knowledge may *not* modulate online speech (Mitterer & Reinisch, 2015).

The limited number of neuroscientific studies investigating this issue are subject to similar confounds. Perre and colleagues (Perre, Pattamadilok, Montant, & Ziegler), for instance, observed that orthographic consistency effects were localised to phonological processing areas such as left BA40 (the surpamarginal gyrus) when participants made lexical decisions. Pattamodilok et al.(Pattamadilok, Knierim, Kawabata Duncan, & Devlin, 2010) also used a lexical decision task and observed that orthographic consistency effects disappeared when phonological processing was interfered with by using repetitive TMS applied to phonological processing areas (the supramarginal gyrus). The same was not observed when TMS was applied to orthographic processing regions. As for the behavioural studies discussed above, there are substantial doubts regarding the extent to which responses in meta-linguistic tasks (such as lexical decision) are a good proxy for normal speech processing (Diependaele, Brysbaert, & Neri, 2012; Gibbs & Van Orden, 1998; Keuleers & Brysbaert, 2011).

One likely explanation for this apparent discord is that the participants in the present investigation were varyingly literate in Devanagari script, which, in contrast to Latin script is not alphabetic – it is an abugida (Daniels, 2020). This raises the intriguing possibility that neural orthography-on-speech processing effects are in fact script-specific rather than universal. It is conceivable that the Devanagari abugida imposes different visual to orthographic mapping requirements in comparison to an alphabetic one as the characters encode consonant-vowel pairs (syllables) rather than sub-syllabic segments (phonemes). Devanagari and Latin scripts’ properties intersect at the conceptual level of sound-symbol mapping and the necessity of assembling sequential symbols to compose words. The evidence presented above suggests that having learned the mappings between orthographic symbols and their phonological renderings *per se* does not necessarily induce changes to the processing of continuous speech in the auditory system as a whole, nor in areas specifically investigated due to the *a priori* evidence of their role in acoustic-phonetic processing of speech.

There is substantial debate about the nature of representation of the acoustic units of human speech sounds. Traditionally it has been assumed that the phoneme is the basic unit of speech, a position that seems “logical” when letters map onto phonemes. However, although it is reasonable to assume some form of phoneme-letter mapping for highly-researched alphabetic writing systems such as English, it is not an account that is likely to be true across the writing systems of the world. Logosyllabic scripts (e.g. Chinese, see Daniels, 2020) have a many-to-one mapping between symbols and sounds, and are fundamentally intransparent (the sound of a word cannot straightforwardly be derived from its symbol), in syllabic scripts (e.g. Kana of Japanese) every symbol encodes a syllable (although there is no transparent relationship between the symbol and the consonant and vowel component), and in abugidas (e.g. Indic and Ethiopic scripts, see Daniels, 2020) every symbol systematically encodes a consonant with a modifier encoding a vowel (or a consonant with an inherent vowel, when unmarked). These different scripts reflect different spoken language units when compared to the phoneme-letter mappings of alphabets (logosyllabaries: morphemes and words, syllabary: syllables, abugidas: mappings at multiple levels of granularity). Indeed, previous research strongly suggests that phoneme units are not ‘needed’ until people learn to read an alphabetic script. A large number of studies has shown that awareness of subsyllabic speech units such as phonemes does not arise spontaneously but that it has to be taught during learning to read (Morais, 2021; Morais, Cary, Alegria, & Bertelson, 1979). Moreover, even in languages with an alphabetic writing system, a phoneme is typically not produced in isolation – speech planning and production take place at either the syllable or segment level (Laganaro, 2019), which map more directly onto syllable-sized orthographic symbols. It is conceivable that learning to map subsyllabic segments to a visual code in alphabetic writing systems might require or induce modifications to auditory processing and representations of speech in order to support the phoneme-level manipulations and representations that are relevant for writing in that specific orthographic system.

We propose that the nature of the speech unit encoded in the orthographic system used by literate individuals must be considered when generating hypotheses about the impact of literacy on speech sound representation and processing. Ultimately, we would argue that literacy in all orthographies is not equivalent, and that drawing conclusions of a universal nature from investigations of alphabetic literacy alone is problematic. As we have previously discussed (Hervais-Adelman et al., 2019), the impact of literacy on visual processing reported by Dehaene and colleagues for alphabetic literates (2010) was not replicated in this group of Hindi-speaking individuals, underscoring the need for further investigations to provide concordant, or discordant, evidence for influential proposals.

An especially intriguing finding of the present study is that the pSTG ROI showed greater functional connectivity with GFMA during spoken sentence processing both cross-sectionally as a function of literacy and longitudinally within-participant as a result of literacy training. The functional connectivity between this region of posterior superior temporal cortex that is associated with acoustic-phonetic processing of phonemes and the handwriting-related areas of the dorsal motor and premotor cortices is of outstanding interest. Literacy is almost never acquired as a purely receptive skill but also involves an important production component when learning to write by hand (but also in typing). It has previously been demonstrated that recognizing (alphabetic) letters activates premotor cortical areas consistent with the representation of the hand habitually used to write (Longcamp et al., 2008). This is compelling evidence for a functional role of graphomotor processes in reading. However, the role of learning to write in developing acoustic-phonetic representations at the level encoded by the script is barely discussed, even though there would be every reason to posit that creating motor-auditory mappings for writing must be as important in becoming literate as learning the visual-auditory bases for decoding script.

While future studies will be necessary to better examine the implications of this functional relationship, the data at hand indicate that, in literate individuals, there is significantly greater coupling between hand-motor regions and auditory processing areas during online sentence processing, in the absence of any orthographic of manual task. Although it is consistent with classical Hebbian processes (Hebb, 1949) that repeated pairings of orthographic tokens with their spoken representations during learning can lead to functional coupling as a result of exercising orthographic output, the relevance of this to spoken language processing is unclear. Importantly, it suggests that we must consider the role of auditory-manual mapping in theories of the role of literacy in the development of phonological representation and processing.

Future, ideally pre-registered longitudinal, studies will be required to systematically examine the potential script specificity (alphabetic vs non-alphabetic) of literacy-induced modulations of responses to speech, in the presence and absence of metalinguistic tasks, and to better understand the role of graphomotor learning in influencing auditory processing of speech.

## Author contributions

Conceptualization: FH, RKM

Methodology: AH-A, FH, RKM

Investigation: UK, RKM, VNT, AG, JPS, FH

Visualization: AH-A

Supervision: FH

Writing: AH-A, FH

## Data and materials availability

Raw data can be made available upon reasonable request to FH. Custom analysis code can be made available upon request to AH-A

